# Atomic force microscopy visualizes mobility of photosynthetic proteins in grana thylakoid membranes

**DOI:** 10.1101/426759

**Authors:** Bibiana Onoa, Shingo Fukuda, Masakazu Iwai, Carlos Bustamante, Krishna K. Niyogi

## Abstract

Thylakoid membranes in chloroplasts contain photosynthetic protein complexes that convert light energy into chemical energy. Photosynthetic protein complexes are considered to undergo structural reorganization to maintain the efficiency of photochemical reactions. A detailed description of the mobility of photosynthetic complexes in real-time is necessary to understand how macromolecular organization of the membrane is altered by environmental fluctuations. Here, we used high-speed atomic force microscopy to visualize and characterize the *in situ* mobility of individual protein complexes in grana thylakoid membranes isolated from *Spinacia oleracea*. Our observations reveal that these membranes can harbor complexes with at least two distinctive classes of mobility. A large fraction of grana membranes contained proteins with quasi-static mobility, exhibiting molecular displacements smaller than 10 nm^2^. In the remaining fraction, the protein mobility is variable with molecular displacements of up to 100 nm^2^. This visualization at high-spatiotemporal resolution enabled us to estimate an average diffusion coefficient of ∼1 nm^2^ s^-1^. Interestingly, both confined and Brownian diffusion models could describe the protein mobility of the second group of membranes. We also provide the first direct evidence of rotational diffusion of photosynthetic complexes. The rotational diffusion of photosynthetic complexes could be an adaptive response to the high protein density in the membrane to guarantee the efficiency of electron transfer reactions. This characterization of the mobility of individual photosynthetic complexes in grana membranes establishes a foundation that could be adapted to study the dynamics of the complexes inside the intact and photosynthetically functional thylakoid membranes to be able to understand its structural responses to diverse environmental fluctuations.

**STATEMENT OF SIGNIFICANCE:** We characterized the dynamics of individual photosynthetic protein complexes in grana thylakoid membranes from *Spinacia oleracea* by high-speed atomic microscopy (HS-AFM). Direct visualization at high spatiotemporal resolution unveils that the mobility of photosynthetic proteins is heterogeneous but governed by the confinement effect imposed by the high protein density in the thylakoid membrane. The photosynthetic complexes display rotational diffusion, which might be a consequence of the crowded environment in the membrane and a mechanism to sustain an efficient electron transfer chain.

## INTRODUCTION

Photosynthesis is a fundamental process that sustains virtually all life on earth. The two photosystems (PSI and PSII), cytochrome *b*_*6*_*f* complex (Cyt *b*_*6*_*f*), and ATP synthase are the major multisubunit membrane protein complexes that catalyze light-driven chemical reactions to generate ATP and NADPH in chloroplasts (1). Light energy is funneled into the reaction centers of each photosystem through light-harvesting complex (LHC) proteins. LHC proteins of PSII (LHCII) are the most abundant membrane proteins in chloroplast thylakoid membranes, and they are also known to play an essential role in photoprotection (2–4). Given the complexity of the light reactions of photosynthesis and their regulation, investigation of thylakoid membrane structure and function has long been a central topic in the field of photosynthesis research.

Thylakoid membranes in plant chloroplasts are organized into intricate structures comprised of highly stacked and non-stacked membrane regions called grana and stroma lamellae, respectively (5, 6). It is well established that PSII and LHCII proteins are predominantly localized in grana, whereas PSI and ATP synthase are exclusively located in stroma lamellae (7). Photo-regulatory processes such as photosynthesis, photoprotection, protein repair, etc. are thought to trigger architectural reorganization in the thylakoids (8). However, the molecular displacement of the photosynthetic complexes responsible for those structural rearrangements is poorly characterized. To investigate the dynamics of photosynthetic proteins in the membrane at high spatial-temporal resolution, a microscopy technique has to meet at least four conditions: (i) The spatial resolution is such that individual complexes can be resolved. (ii) The observation is performed in physiological conditions. (iii) The temporal resolution is high enough to track molecular displacements. (iv) The detection method is not invasive to preserve native-like organizations.

Electron microscopy (EM) meets the first and fourth conditions, indeed, EM studies have shown that grana are highly packed with membrane proteins, where PSII and LHCII form a protein supercomplex (9–11). It has also been suggested that the macroorganization of PSII-LHCII supercomplexes plays a critical role during photoprotection, which implies a high significance of their diffusion rates within the thylakoid membrane (12–14). Optical microscopy based on fluorescence detection, on the other hand, fulfills conditions (ii) and (iii). By following chlorophyll (Chl) fluorescence emitted from photosynthetic membrane proteins, it is possible to infer dynamics of thylakoid membranes at high temporal resolution. However, this strategy remains blind to photosynthetic proteins themselves, which provide the binding sites for Chl. In spite of the efforts to improve the spatial resolution using super-resolution optical microscopy (15–18), the information obtained is still somewhat indirect due to the high abundance of Chl pigments in the membrane which limits the specific assignment of the actual photosynthetic complex.

Atomic force microscopy (AFM) is a unique technique that enables direct observation of macromolecules at high spatial resolution (XY < 1 nm and Z < 0.1 nm) in aqueous conditions. AFM has been used to characterize the structure and organization of thylakoid membranes (19–24). These studies support and complement EM reports about how PSII and LHCII organization is affected by illumination (20, 22, 23). Despite the scope of AFM in the characterization of the thylakoid membrane, studying its dynamic behavior was elusive because of the AFM’s intrinsic low imaging speed. Thanks to the development of high-speed AFM (HS-AFM) (25–29), it is now possible to visualize real-time dynamics of biological macromolecules. Thus, HS-AFM has the potential to meet all four conditions stated above. Indeed, HS-AFM has been applied to study dynamic events of purified molecules (30–34), membrane proteins embedded in lipid bilayers (35, 36), and even biological tissues (37–39). Here, we employed HS-AFM to characterize the *in situ* dynamic behavior of the photosynthetic protein complexes in grana thylakoid membranes isolated from *Spinacia oleracea*. Our HS-AFM observations indicated that the fraction of mobile membrane proteins is less than 10% of the total population. The high temporal resolution permitted herein showed that the diffusion of the protein complexes is mainly rotational. Our results also indicated that the diffusion of the photosynthetic membrane proteins is heterogeneous not only between different grana membranes, but also within a single granum. This characterization provides a foundation for studies of thylakoid structural response to different environments.

## MATERIALS AND METHODS

### Grana sample preparation

Grana membranes were prepared from spinach (*Spinacia oleracea*) according to the previous method (40), except for the following modification. Spinach leaves were obtained from a local store and kept in the dark overnight at 4 °C. Digitonin (the final concentration at 0.7% w/v; MilliporeSigma, St. Louis, MO) was used to solubilize chloroplasts (0.4 mg Chl/mL) at 4 °C for 30 min in the buffer containing 50 mM phosphate (pH 7.2), 300 mM sucrose, and 10 mM KCl. Crude grana fractions were removed by centrifugation at 1,000 × g for 3 min at 4 °C. The supernatant was centrifuged at 1,000 × g for 5 min at 4 °C to sediment taller grana. The supernatant was further centrifuged at 1,000 × g for 10 min at 4 °C. The pellet containing shorter-height grana was resuspended in the same buffer and immediately used for AFM observation. Different preparations of grana membranes were considered to be biological replicates. Repeated measurements on the same grana preparation were considered to be technical replicates. Prior to initiating HS-AFM experiments, several preparations (∼6) of grana membranes were routinely inspected by conventional AFM to assure reproducibility and uniformity in appearance and dispersion of individual grana discs.

To determine the composition of the membrane, the samples were solubilized with standard Laemmli sample buffer and separated by electrophoresis using sodium dodecyl sulfate polyacrylamide gel prepared using Any kD TGX precast protein gels (Bio-Rad, Hercules, California). Separated proteins in gel were electroblotted onto a polyvinylidene difluoride membrane using Trans-Blot Turbo transfer system according to the manufacturer’s instruction (Bio-Rad, Hercules, California). Primary antibodies specific for D1 (PSII), PsaD (PSI), Lhcb2 (LHCII), Cyt *f*, and AtpB were obtained commercially (Agrisera, Vännäs, Sweden) and used according to their recommendations.

### Atomic force microscopy

To assess the quality of the solubilized membranes, grana membranes were deposited on freshly cleaved mica in high ionic strength buffer (10 mM Tris-HCl, pH 8.0, 150 mM KCl, and 25 mM MgCl_2_) (41) and incubated at room temperature for 1–3 h. Mica was rinsed with water ten times and dried under N_2_ gas flow for 2 min. We used a Multimode AFM Nanoscope V (Bruker Co. California) and performed the observation as described previously (22).

Grana membranes were diluted 5-to 10-fold in high ionic strength adsorption buffer. Two microliters of the diluted sample were deposited on freshly cleaved mica and incubated for 1 h in the dark. Weakly bound membranes were removed by rinsing 10 times with imaging buffer (10 mM Tris-HCl, pH 8.0, 5 mM MgCl_2_, 150 mM KCl) followed by a gentle, brief (2 s) puff of high purity Argon gas. The sample was immediately immersed in 2 µL of imaging buffer. For scanning, the HS-AFM bath contained the same imaging buffer. We used the Ando-model HS-AFM (Research Institute of Biomolecule Metrology, Tsukuba, Japan) (28) equipped with a near-infrared laser (830 nm wavelength) to minimize chlorophyll excitation during observation. All optical components were adjusted to near infrared region except for an objective lens. We used a wide area scanner (maximum range: 6 × 6 μm^2^ in XY-directions and 1 μm in Z-direction). First, we set the scan range between 1 × 1 μm^2^ to 4 × 4 μm^2^ in order to find appropriate membranes. Then, we moved the stage to place the membrane at the cantilever position and observed it with scan range of 150∼500 nm^2^ at 1 frame s^-1^. The samples were scanned in liquid using tapping mode at room temperature. The deflection of micron-sized cantilever (AC10DS, Olympus, Japan, spring constant ∼0.1 N/m, resonance frequency 400∼500 kHz in liquid) was detected using an optical beam detector. The free-oscillation amplitude of the cantilever (A_0_) was set to ∼2 nm, and the set point (A_s_) of feedback amplitude was set to about 0.9A_0_. We optimized data acquisition (e.g. low amplitudes and high A_s_/A_0_ ratio were chosen to achieve the highest resolution at the lowest force in order to preserve the membrane structure while minimizing artifacts introduced by tip-sample interactions) and data analyses to deconvolute the noise intrinsic to our HS-AFM (see below) to minimize over-interpretation and maximize unbiased observations. The detailed procedure of HS-AFM observation was described elsewhere (42).

### Data analysis

Individual frames from HS-AFM movies selected from a second biological replicate experiment were processed using customized algorithms written in Igor Pro (Wave Metrics Inc. Oregon). First, noise was reduced by Gaussian filtering followed by a flattening filter to accurately measure heights. Second, entire patches were tracked using a 2D correlation method to correct and minimize lateral drift (31). Briefly, a region of interest (ROI) in a first frame is chosen as reference for aligning subsequent frames using 2D cross correlation:

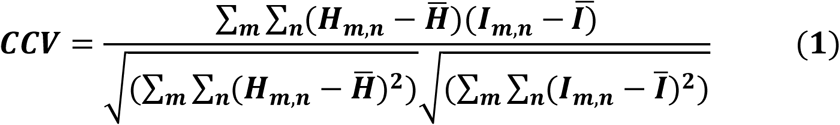

where H and I denote the height values at a pixel point (m, n) for the targeted ROI at different time points and the initial one (from the first frame), respectively. 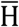 and Ī are height mean values of the matrix. The region of the maximum CCV value found defines the corrected origin of the image. Particle dimensions (median heights, diameters, center of mass, etc.) were obtained from those selected by thresholding segmentation (package features) from the particle and pore analysis module included in SPIP(tm) (Hørsholm, Denmark). Dimensional fluctuations and spatial displacements were tracked, plotted, and fit using customized scripts written in Wolfram Mathematica® (Illinois) or Igor Pro. The goodness of fit for normal distributions was done using the Akaike information criterion. The contrast of high-resolution images was digitally adjusted to facilitate the visual detection of dimeric structures chosen for particle analysis; therefore, small and membrane-embedded proteins appear invisible. Particle MSD was calculated according to:

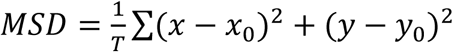

where *x* and *y* are the particle’s center of mass coordinates (center for those points inside a particle with Z values above the mean contour height) at different time points; *x*_0_ and *y*_0_ represents the initial *x,y* center of mass coordinate; *T* = total duration of observation. To assure that the mobility of the photosynthetic complexes is distinguishable from the intrinsic noise of the scanning, we compare the MSDs obtained for dimeric complexes inside the membrane to those of particles of similar dimensions (∼20 nm in diameter) directly attached to the mica substrate. The MSD trajectories of those fixed particles are virtually flat over a period of 80s and with low values (less than 2nm^2^) as shown in Fig. S1, in contrast to those obtained for complexes in the grana membranes.

The MSD traces obtained from complexes in the membranes were then fit with two diffusion models: confined (Eq. 2) (43, 44) and Brownian (Eq. 3).

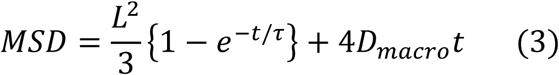

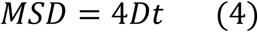

where *D*_*macro*_ = macroscopic diffusion coefficient, *L* = confined domain, τ = equilibration time, *D* = the diffusion coefficient of natural diffusion and *t* = time interval. The microscopic diffusion coefficient (*D*_*micro*_) in the confined diffusion model can be obtained from Eq. 4 (43, 44).

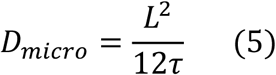

To establish whether or not the particle’s diffusion properties were identical, we determined the relationship between L and the initial diffusion coefficient (D_initial_ at *≈* 4 s) or D_micro_ obtained from Eq. 4. We confirmed an apparent particle’s segregation from this correlation by applying the k-means clustering criteria. The diffusion coefficients reported in this study were obtained from the best fit (Brownian or confined) to the average trace resulted from each subgroup. The goodness of the fitting was evaluated by determining the squared correlation coefficient R^2^.

To dissect the molecular rotational movement, we calculated the correlation coefficient of 2D image (31). After tracking a selected molecule to eliminate lateral diffusion effects, we defined it into a rectangular ROI to calculate the 2D correlation coefficient frame by frame with Eq. 1.

## RESULTS

### HS-AFM visualizes dimeric photosynthetic complexes without altering macromolecular organization in grana membranes

We isolated grana membranes from spinach using digitonin as described previously (40) (Fig. S2). This preparation yielded small stacks of grana discs with high content of PSII, LHCII and Cyt b_6_f complexes as previously shown (21). We optimized the HS-AFM setup for imaging photosynthetic proteins in grana membranes, such that deflection of the AFM cantilever was detected using a near-infrared laser (830 nm wavelength) to minimize Chl excitation. The membranes were scanned at one frame per second in buffer solution.

HS-AFM observations indicated that macroorganization of grana membranes and associated protein structures were well preserved (Figs 1 and S4). As shown previously (19, 22, 24, 45), grana membranes were packed with dimeric complexes with an overall density of ∼1456 ± 230 particles/µm^2^. Dimeric structures were distributed throughout the membranes, but their structural arrangement appeared to be disordered (i.e. randomly oriented throughout the membrane with no crystalline arrangements). We observed a bimodal distribution of height of the dimeric structures, which is consistent with previous results (Fig. S3, *a* and *b*) (21). Immunoblot analysis detected the existence of PSII and Cyt *b*_6_*f* in our grana membranes prepared using digitonin (21) (Fig. S2 *a*). Certainly, the unambiguous distinction of PSII and Cyt *b*_6_*f* is highly desirable whenever both of them are present in the membranes. However, imaging by itself is insufficient to discriminate these two dimers. Comparison of available structures of Cyt *b*_6_*f* and PSII indicates that their dimensions are very similar (46, 47). Both Cyt *b*_6_*f* and PSII have similar shapes and lateral sizes of ∼16 and ∼20 nm, respectively. Moreover, in both cases, their membrane lumenal protrusions are separated by ∼8 nm and their expected heights differ by only ∼1 nm (21). These characteristics challenge an unambiguous discrimination of each individual complex but are sufficient to estimate the presence of distinct populations of complexes. There is a structural report that succeeded in distinguishing both dimers by affinity-mapping AFM (21), which specifically recognizes plastocyanin-Cyt *b*_*6*_*f* interactions. It is worth noting that we did not observe ATP synthase by HS-AFM in our grana membranes. It is known that solubilization by digitonin cannot remove ATP synthase completely from the grana preparation, as observed in our immunoblot analysis (Fig. S2 *a*) and a previous study (21). For the purpose of this study, it was more important to prepare the membrane samples closer to native conditions than to remove ATP synthase completely by using stronger detergents. In fact, the ATP synthase content in our grana membranes was reduced 2-fold from leaves, which was sufficient for our observation. For HS-AFM observation, the samples were 5-to-10-fold further diluted to adjust their density on the surface, such that individual grana discs could be isolated. Furthermore, we avoided scanning membranes with large protruding structures, and thus we minimized the probability to study membranes containing ATP synthase complexes.

**FIGURE 1.**
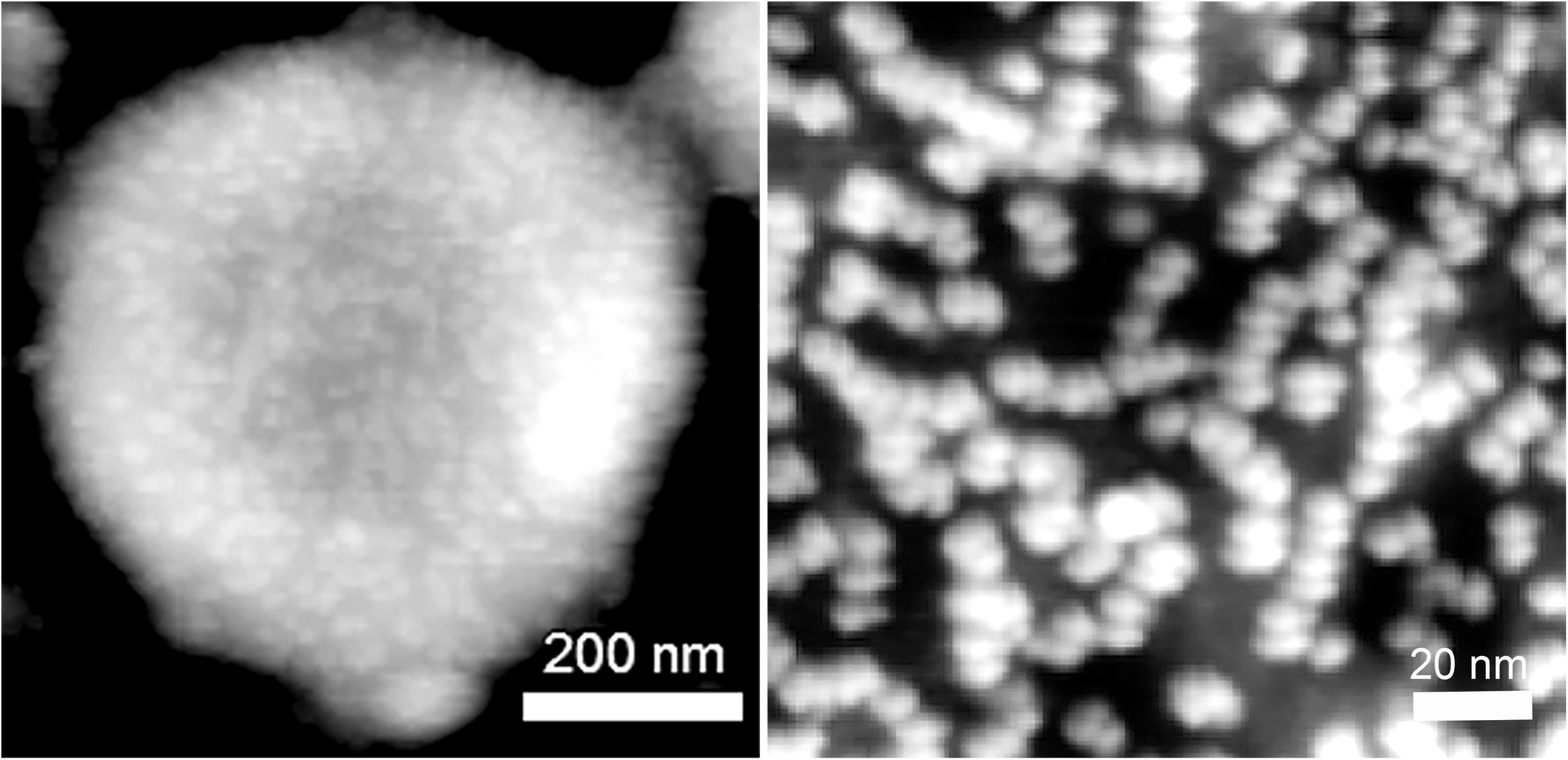
HS-AFM micrographs of the representative grana membrane. (*a*) HS-AFM image of a grana membrane. Z-scale, 95 nm. (*b*) Magnified averaged HS-AFM image of a grana membrane. Z-scale, 6.0 nm. The image was acquired at 1 frame per second and averaged over 50 frames. The contrast of the membrane was adjusted to enhance the dimeric structures; small, membrane -embedded proteins such as LHCII are invisible.

We also calculated the nearest neighbor distance (NND) distribution function (Fig. S3 *c*). A main peak centered at ∼20 nm flanked by shorter (∼16 nm) and longer (∼25 nm) distances was well fitted, which is also consistent with previous results (22, 48). These results observed by HS-AFM are qualitatively comparable with those observed by conventional AFM performed in air (Fig. S2, *b*-*d*). The protein density and NND distribution observed by HS-AFM are also consistent with those observed by EM (e.g. the samples with no light treatment (45, 49). Taken together, these results demonstrated that HS-AFM is a suitable technique to image the thylakoid protein complexes at high spatial resolution *in situ* without altering their intrinsic macroorganization.

### HS-AFM revealed heterogeneous protein diffusion in individual grana membranes

To analyze the dynamics of photosynthetic complexes in grana membranes, we performed HS-AFM observation for 60 s or more per sample. HS-AFM images of representative grana discs are shown in Fig. S4 *e*. HS-AFM was performed on a total of 19 grana discs from two biological replicate preparations. We tracked individual protruding dimeric structures to calculate the mean square displacement (MSD, Eq. 1). Based on the level of lateral displacement, grana membranes that we observed here could be divided into two groups. The first group, which comprises approximately 90% of the total grana membranes observed in this study, was termed QSM membranes because they contained dimeric structures with <u>q</u>uasi-<u>s</u>tatic <u>m</u>obility (Fig. 2 and Video S1). The distinct dimeric structure of each particle in a QSM membrane was still apparent after averaging 50 frames of the HS-AFM images (Fig.2 *a*), which indicates that the lateral displacement was confined to a few nanometers. The MSD values of 53 dimeric structures in this representative QSM membrane show that the molecular displacement was less than 10 nm^2^ (Fig. 2 *b,c*). These values are slightly above the displacement of free particles directly fixed onto the substrate (Fig. S1). Therefore, these displacements are, in part, due to the residual mechanical drift in HS-AFM apparatus that our algorithms did not eliminate. However, histogram of the MSD values at 50 s depicts the distribution (∼4 nm^2^) greater than the mechanical drift (Fig. 2 c). It is also worth stressing that the photosynthetic complexes tracked in our samples are never in direct contact with the substrate (Fig. S4). Our solubilization method yields grana patches with two or more appressed grana discs as indicated by their height profiles 60 ± 8.0 nm, (Fig S7). AFM can only probe the mobility of the complexes protruding from the top layers, which are at least 50 nm (about 4 lipid bilayers or 2 grana discs) above the mica surface, therefore these complexes most likely interact with the lipid bilayer underneath and/or other thylakoid complexes. Thus, we conclude that the dynamic behavior of the complexes reported in this study is independent from the membrane-substrate interactions. Lastly, the average MSD trace of all structures (thick line in Fig. 2 3 *bc* insect) was well fitted to a confined diffusion model (Eq. 2). Because the grana membranes used in this study showed preserved macroorganization (expected protein density, dimensions, and NNDs) as demonstrated in the previous section (Figs 1, S2 and S3), we consider that the quasi-static mobility observed in the QSM group does not indicate an aberrant state of the membranes. In the second group, which comprises about 10% of the total grana membranes observed in this study, most dimeric structures showed quasi-static mobility and sometimes appeared to be clustered (arrowheads in Fig. S5). However, there was a subpopulation of dimeric structures that displayed larger displacements. Fig. 3 *a* shows representative time-lapse HS-AFM images revealing such dimeric structures (see also Video S2).

**FIGURE 2.**
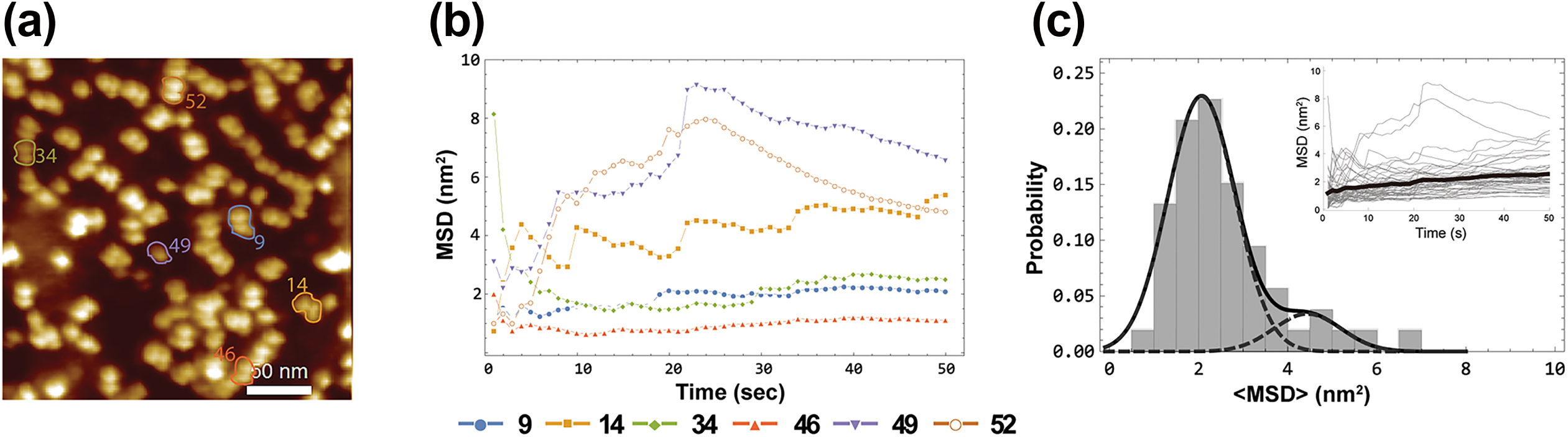
HS-AFM characterization of the protein mobility observed in the first (QSM) group of membranes. (*a*) Time-average image of a representative HS-AFM movie. Each image was acquired at 1 frame per second and averaged over 50 frames. (*b*) Mean square displacement (MSD) trajectories of selected proteins with different surroundings; (*c*) Probability distribution of the average MSD of 53 proteins over 50 sec detected in the entire membrane. Inset shows all 53 MSD trajectories (gray lines) and the average trajectory of all proteins (superimposed black thick line). The confined diffusion model agrees well with the average displacement of proteins in the QSM group of membranes.

**FIGURE 3.**
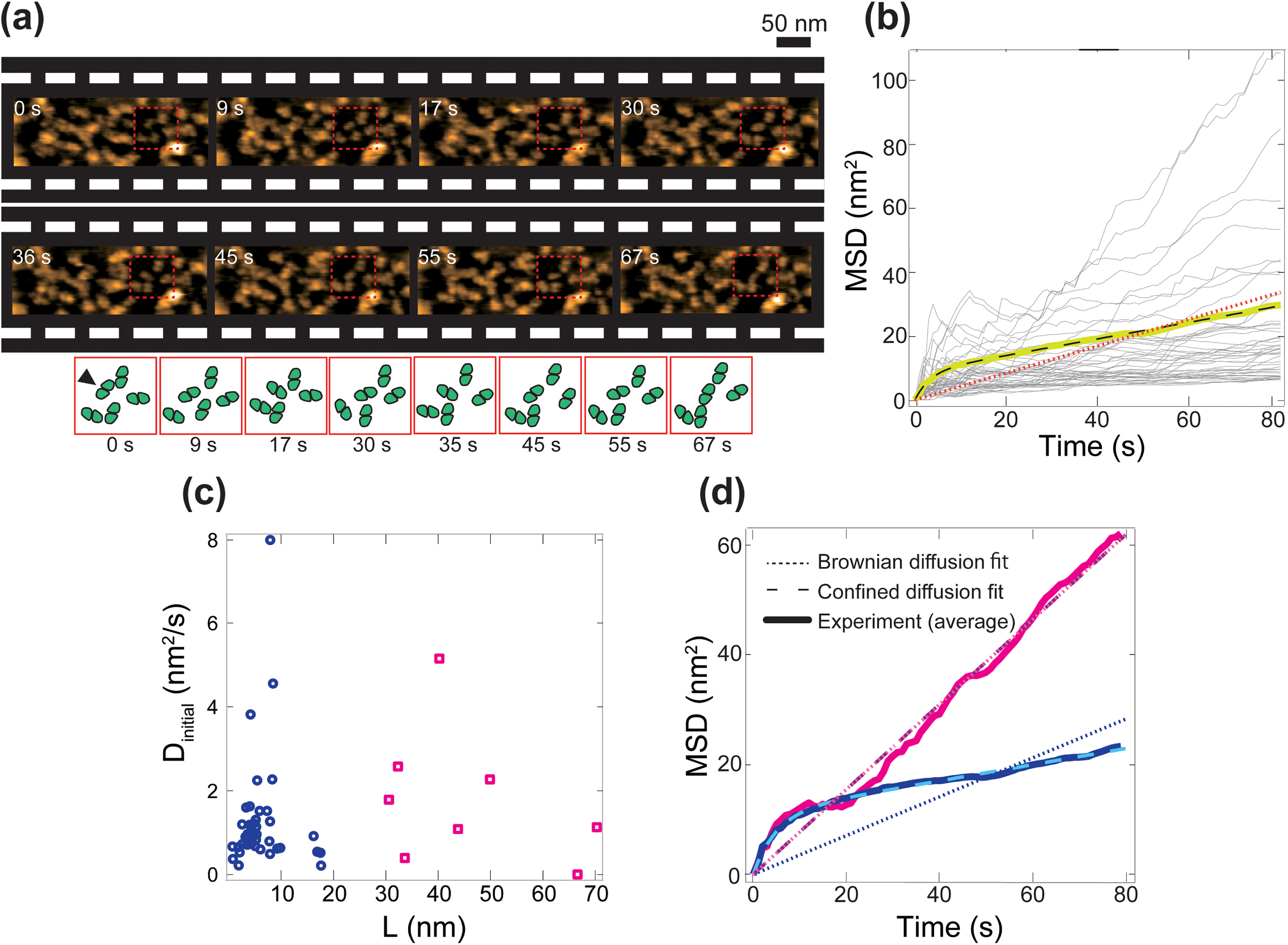
HS-AFM characterization of the protein mobility observed in the second (VPM) group of membranes. (*a*) Time-lapse HS-AFM images showing the heterogeneous protein mobility in the VPM group of membranes. Images were acquired at 1 frame per second. Corresponding illustrations of few proteins enclosed in the red squares are presented to facilitate the visualization of the protein dynamics. The particles in the cartoon were drawn freehand by visually tracing the protein’s contour in the HS-AFM images. Z-scale, 7.0 nm. (*b*) MSD trajectories of the proteins selected in this membrane (N = 50). Gray lines are individual proteins’ MSD trajectories; superimposed yellow line is the MSD average of all proteins; dashed black line is a fit to a confined diffusion model (Eq. 2); and orange dotted line is a fit to the Brownian diffusion model (Eq. 3). (*c*) Relationship between each protein’s confined length and its initial diffusion coefficient (4 s). A protein’s mobility can be segregated into two distinct groups as shown in blue circles and magenta squares, according to the k-means criteria. (*d*) Average MSD traces of two distinct groups (blue and magenta in *c*); dotted and dashed lines are the fits to Brownian and confined diffusion models, respectively.

The individual MSD traces were variable among the selected dimeric structures (thin gray lines), and some of them showed larger displacements. (Fig. 3 *b*). Therefore, the second group of grana membranes was termed VPM for <u>v</u>ariable <u>p</u>rotein <u>m</u>obility. The average MSD trace of all observed dimeric structures in the VPM membranes (yellow line in Fig. 3 *b*) fitted well to a confined diffusion model (Eq. 2, black dashed line, R^2^ = 0.996) as compared to a Brownian diffusion model (Eq. 3, orange dotted line, R^2^ = 0.981). Using Eqs. 2 and 4, we obtained the average diffusion coefficient of ∼1 nm^2^ s^-1^ that was consistent with previous Monte Carlo simulation reports (see Discussion). This result indicates that, overall, the dimeric structures in the VPM grana membranes still exhibited dynamics that can be explained by a confined diffusion model, similar to those observed in the QSM group.

The average MSD trace of the entire population of particles in the VPM membranes may underestimate a certain subpopulation, which apparently showed an unconfined diffusion. To carefully determine the diffusion of each individual dimeric structure, we first fitted each MSD trace to the confined diffusion model. Next, we extracted each confined domain (L) and correlated it with the diffusion coefficient during the first 4 s (D_initial_). We then applied k-means clustering criteria (50). The results showed that most of the dimeric structures (84%) (blue circles in Fig. 3 *c*) were each confined to a small region (average L = 5.8 ± 0.04 nm). In contrast, the remaining population (magenta squares in Fig. 3 *d*) displayed larger displacements (average L = 44.3 ± 17.6 nm). We re-calculated the average MSD trace separately for these two populations. The average MSD trace (blue trace in Fig. 3 *d*) from the larger population was better fitted to a confined diffusion model (light blue dashed line; Eq. 2, R^2^ = 0.997) than to a Brownian diffusion model (blue dotted line; Eq. 3 R^2^ = 0.939). On the other hand, the fitting of the average MSD trace from the smaller population (magenta trace in Fig. 3 d) is more ambiguous. Either a confined (light pink dashed line; Eq. 2, R^2^ = 0.983) or Brownian diffusion model (magenta dotted line; Eq. 3, R^2^ = 0.983) fits the average MSD trajectory in similar fashion, but neither fit is as well-adjusted as observed for the larger confined population. We interpret this characteristic as an indication that this type of membrane harbors a fraction of complexes with the potential of being unconfined and adopt a Brownian diffusion behavior. In other words, we propose that these membranes can allow the coexistence of two populations of complexes: a large fraction of proteins mainly confined and another one that either escape or are at the verge of escaping (note the curvature of the average MSD during the first 20 seconds) and diffuse freely and faster. The reason for this behavior is unknown to us at this time. We thus refer to this phenomenon as heterogeneous mobility. Note that the tip-sample interactions do not affect the detected proteins mobility. We did not detect any bias of the tracking trajectories to the fast-axis (X-axis) as shown in Fig. S6, therefore, we conclude that the external force applied by the AFM tip on the membrane could also be safely neglected. In an attempt to explain the nature of this heterogeneous mobility and/or to distinguish PSII from Cyt *b*_*6*_*f* by their mobility, we interrogated if there was any correlation of the complexes’ mobility and their height (i.e. smaller complexes might diffuse faster than taller/larger ones). We measured the height of dimeric proteins in the VPM membranes and correlated them to their MSD. There was no correlation between complexes’ heights and MSD (Fig. S8). From this result, we consider that the complexes’ mobility was inconclusive to distinguish whether the mobile fraction corresponded specifically to PSII or Cyt *b*_*6*_*f*. We also noticed that the protein density (1594 ± 364 particles/μm^2^) in the VPM membranes was not statistically different from that in QSMs (P = 0.7, Student’s *t-*test). Interestingly, the median height distribution of the particles in VPM membranes was broader compared to that of QSMs, and it was best fitted with a single Gaussian based on Akaike information criterion (Fig. S9 *a*). Unfortunately, these membranes constitute a small fraction (only 10% of the membranes), and the identification of individual complexes is hindered due to their clustering, which prevents us from establishing a reliable statistical significance of this observation. We also detected fluctuations of the NND throughout the observation (i.e. the NND at time 0 s varies continuously), which could be expected if indeed there is a more mobile population of complexes. These fluctuations of the NND over time limit a quantitative comparison (a particular time must be chosen arbitrary) to similar distributions from the QSMs. Nevertheless, by comparing the initial time (0 s) of the observations, it appears that the NND in these membranes is modestly shifted towards shorter distances compared to those of QSMs (Fig. S9 b). This would be consistent with the presence of protein clusters. Protein clustering might favor molecular unconfinement by creating unpopulated domains within the membrane. Furthermore, we detected modest fluctuations of the complexes’ height throughout the observation. We compared the heights and NNDs of the particles (first and last frame) in the same VPM membrane. There are visible differences between both height and NND distributions (Fig S9 *c* and *d*), suggesting that the global organization of the membrane, as well as the level of protrusion of the complexes, were slightly changed during our observation. It is worth mentioning that we occasionally (less than 1% of the total number of membranes) observed grana membranes that contained larger populations of mobile dimeric structures (>80%) than immobile ones (Video S3). In such cases, dimeric structures tended to collide with each other more frequently. Due to the scarcity of these membranes, we were unable to perform any type of statistical analysis to characterize their structural organization and/or mobility properly. The origin of this behavior is also unknown at this time, but it could indicate a different state—including an anomalous one—of the membrane induced by some external factor.

### HS-AFM visualized rotational displacement of dimeric structures in grana

High temporal resolution of HS-AFM in this study also enabled us to analyze rotational displacement of the dimeric structures. In the VPM grana membranes, which contained the dimeric structures with heterogeneous diffusion, we measured changes in the angle (θ) of two adjacent structures during a 1-min observation period (magenta and blue arrows in Fig. 4 *a* and Video S4). The results indicated that the angle changed from 20 to 80 degrees within a few seconds (Fig. 4 *b*). To further illustrate the rotational diffusion observed here, we calculated a 2D-correlation coefficient of variation (CCV; Eq. 5) of the dimeric structures as shown in Fig. 4 *c*. The dimeric structure showing rotational diffusion exhibited constant fluctuations with an average CCV value of 0.65 throughout the observation (red trace in Fig. 4 *c*). By averaging the total time-lapse images, the dimeric morphology disappeared because of the rotational displacement (red profile in Fig. 4 *d*). In contrast, the other dimeric structure showed an average CCV value of 0.83 (blue trace in Fig. 4 *c*), and the dimeric morphology of this structure stayed the same after averaging the time-lapse images (blue profile in Fig. 4 *d*).

**FIGURE 4.**
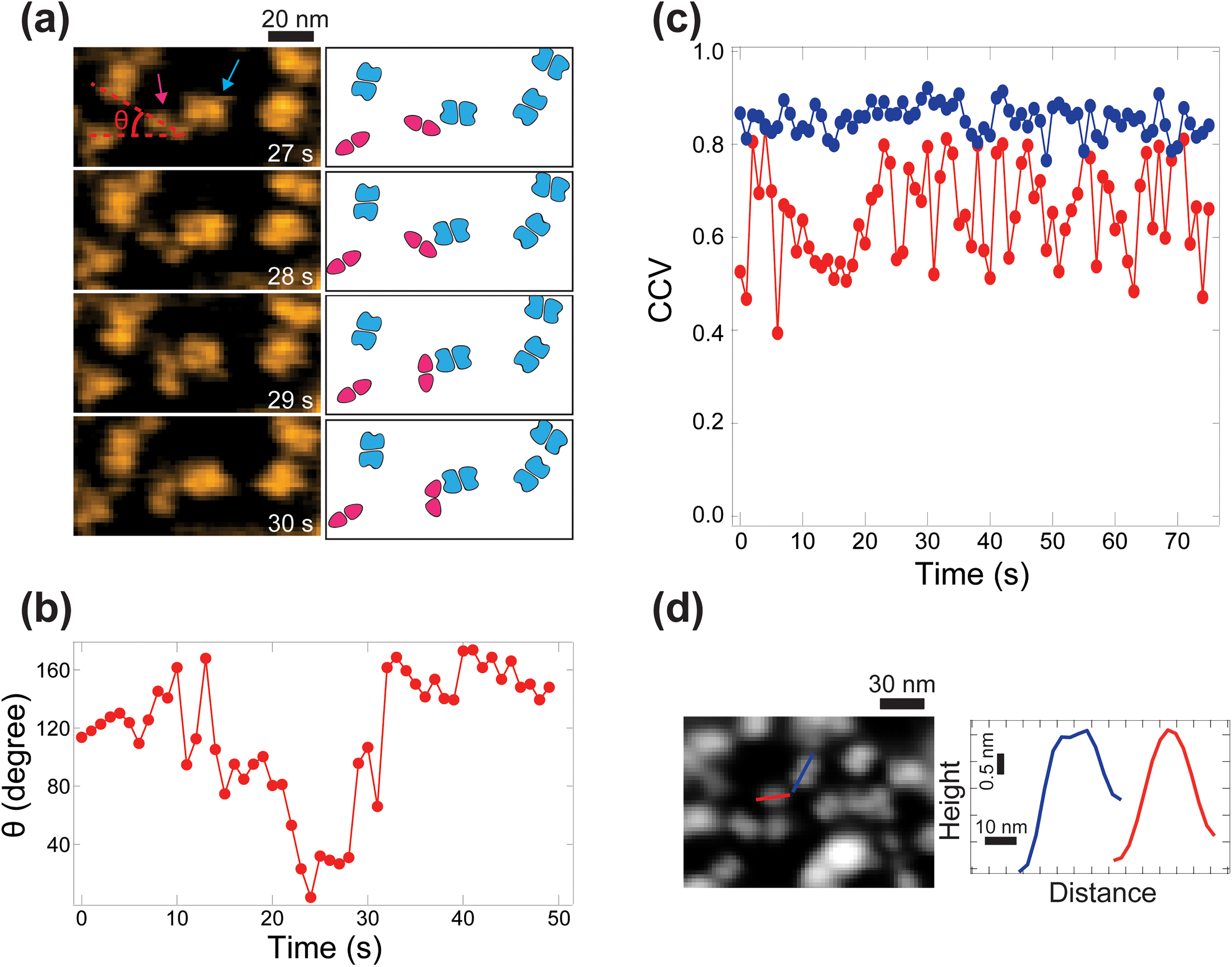
HS-AFM characterization of the rotational displacement of a dimeric structure. (*a*) Time-lapse HS-AFM images illustrating the rotation of a dimeric protein (magenta arrow) with respect to its neighbor protein (blue arrow) captured at 1 fps. Corresponding illustrations are shown in the right (the particles were drawn by following the protein’s contour as described in Fig. 3). Z-scale, 7.0 nm. (*b*) Variation of the angle (θ) in panel *a* over 50 s. (*c*) Time-course examples of the 2D correlation coefficient of variation (CCV) for two different proteins. Non-rotational symmetry will produce CCV values closer to 1. Rotational displacement is depicted by the red trace. Negligible rotational displacement is illustrated by the blue trace. (*d*) Left, the average HS-AFM image. Right, height profiles indicated in averaged AFM images (left).

It is known that PSII-LHCII supercomplexes in grana can be organized into a higher order of associations (e.g. megacomplexes or two-dimensional crystalline arrays (51). Intriguingly, it has been suggested that there are two types of PSII megacomplexes, in which 80% of the supercomplexes show parallel associations, whereas the other 20% interact in a non-parallel manner with variable associations between the two supercomplexes (52). The rotational displacement of the dimeric structures detected here is in line with the idea that PSII-LHCII supercomplexes might associate with their neighbors at slightly different orientations.

## DISCUSSION

This contribution aims to establish that HS-AFM is a powerful technique to study the dynamic behavior of photosynthetic protein complexes in plants under biologically relevant conditions. Indeed, we were able to characterize the mobility of individual complexes in highly crowded grana membranes. Despite extensive previous investigation, there is currently a lack of consensus about the actual diffusion properties of photosynthetic complexes in thylakoids. This is, in part, because the experimental results are indirectly obtained from the total pool of Chl-binding proteins in the membranes, and because the large size of the supercomplexes limits the scope of molecular simulations (8, 12, 53–57). Our HS-AFM observations provide insight into diffusion of photosynthetic proteins in grana thylakoid membranes, which is essential information for understanding relevant biological processes such as photosynthesis, photoprotection, photodamage, and repair.

The HS-AFM scanning of grana membranes in solution at a rate of one frame per second was sufficient to resolve the dimeric structure of the photosynthetic protein complexes (PSII or Cyt *b*_*6*_*f*). In our preparation, the diameter of grana discs varied from 250 to 600 nm, so in order to observe the entire disc, the scanning area was set at 300-500 nm^2^ (∼ 150×150 pixels). However, higher temporal resolution can be achieved by scanning smaller areas (28).

Our HS-AFM observation revealed the presence of at least two groups of isolated grana membranes (QSM and VPM) according to the diffusion behavior of observed dimeric protein structures. QSM represents the majority of grana membranes observed (∼90% of ∼19 grana discs) and contains dimeric structures with quasi-static mobility that fits a confined diffusion model (Fig. 2). As the remaining 10%, VPM contains dimeric structures showing larger displacements and a higher diffusion rate than the first group. The diffusion model for VPM appears to be fitted with both confined and Brownian models, but there are individual trajectories that fit only a Brownian model because of their high diffusion rates (Fig. 3b), suggesting that those complexes are no longer confined. The architecture of both types of membranes are qualitatively similar (lack of crystalline arrays and similar complex densities), nevertheless it appears that the global organization is slightly different (i.e., more protein clustering). We do not have evidence to suggest the exact reason for this heterogeneity. As we observed a similar density of dimeric structures in both QSM and VPM grana membranes but shorter NNDs, possible reasons for this heterogeneity could be different ratio of LHCII to PSII and/or different lipid compositions in the membranes, which might originate from different parts of thylakoid membranes. Also, it has been previously shown that, under fluctuating light, the organization of PSII-LHCII supercomplexes could undergo reversible transitions from crystalline to fluid phases (22, 45). Therefore, we speculate that the heterogeneous protein mobility observed in this study might partially reflect different physiological conditions or transient events related to photoacclimation mechanisms in the leaves from which the grana were isolated. Future experiments using plants acclimated to different light environments will provide a detailed connection between our HS-AFM observation and physiological mechanisms. In this study, we provide a proof of concept that HS-AFM is a suitable technique to study the protein dynamics in thylakoid membranes.

Molecular confinement is well-described in highly crowded membranes such as thylakoid membranes (55), which would have a significant impact on protein mobility (e.g. diffusion paths and velocities). Fluorescence recovery after photobleaching (FRAP) experiments both in intact or broken chloroplasts and isolated grana membranes have shown that about 80% of chlorophyll-binding proteins are immobile (12, 56). HS-AFM allows us to visualize the displacement of individual dimeric photosynthetic complexes in thylakoid membranes and to characterize their diffusion. Our HS-AFM observation indicated that 90% of observed grana contain immobile proteins, similar to the results reported by FRAP experiments. This suggests that the overall protein immobility in chloroplasts observed by FRAP might reflect the confined protein mobility occurring in grana.

Additionally, our HS-AFM observation revealed that protein diffusion can be segregated even within the same grana disc (Figs 3 and 4). It has been shown by using *in vitro* reconstituted lipid membranes with a protein surface density of ∼50% that local protein density is correlated with protein mobility, demonstrating that molecular confinement has a significant effect on protein diffusion in membranes (35). The grana thylakoid is however a special type of membrane expected to restrict diffusion of its proteins more than plasma membranes as it is highly packed with a ratio of protein:lipid of 86:14 (w/w) (1). AFM scanning can only visualize those domains protruding from the lipid bilayer (e.g. the oxygen evolving complex in PSII). In the thylakoid membrane, there are many more proteins embedded in the lipid bilayer which cannot be detected by AFM. These lipid-embedded proteins could be hidden inside the lipid space between the protruding bright particles, which appear darker in the micrographs due to their lower heights (z-values). These embedded proteins could also affect the mobility of the visible ones. LHCII proteins are most abundant in grana and are suggested to be both immobile and mobile as they can associate with PSII and interact with other LHCII proteins, reorganizing different protein complexes in response to light fluctuations (8). The heterogeneity of protein mobility in a single granum observed here might indicate such different situations of LHCII, some of which are strongly associated with PSII, whereas others diffuse freely between PSII supercomplexes and thereby affect the apparent mobility of dimeric structures. To fully understand diffusion of dimeric proteins (PSII and Cyt *b*_*6*_*f*) in thylakoid membranes, it will be necessary to consider the effect of these embedded membrane proteins.

From our HS-AFM observation and analysis, we calculated that the diffusion coefficient of dimeric structures in grana is approximately 1 nm^2^ s^-1^ eq. (4). The results from FRAP measurements estimated a diffusion coefficient of ∼100 nm^2^ s^-1^ (56), 100-fold higher than our observation, which is most likely due to the fact that our HS-AFM tracks individual dimeric structures in grana, whereas FRAP measures the ensemble of Chl-binding proteins. Coarse-grained simulations of individual PSII complexes calculated a diffusion coefficient of 100,000 nm^2^ s^-1^ (57). However, this simulation did not account for the molecular crowding effect. Monte Carlo simulations based on FRAP experimental data that include the effect of molecular crowding calculated a diffusion coefficient of 1 nm^2^ s^-1^ (48), which agrees well with our direct observation of these complexes. These results emphasize that it is essential for understanding diffusion of membrane proteins to measure not only the mobility of individual molecules but also to consider the effect of molecular crowding (55, 56), which is only achievable experimentally by HS-AFM.

HS-AFM imaging of thylakoid membranes also provided direct evidence of rotational diffusion of photosynthetic complexes (Fig. 4). Rotational mobility of membrane proteins have been detected by many different techniques, including HS-AFM (35, 58). The rotational diffusion is a consequence of combined effects: the protein’s structure, its location, its function, protein-protein and protein-lipid interactions, as well as the local environment (59–62). In a much more simplified, reconstituted membrane (50% of protein density and ∼10-fold smaller than PSII reaction core), Casuso *et al*. were also able to detect by HS-AFM the rotational motion of OmpF homotrimers (35). In both cases, photosynthetic and reconstituted membranes, the rotation coefficient of the proteins could not be determined, because of the heterogeneous mobility of the proteins due to the confinement effect. It is conceivable that during the photosynthetic reactions, protein rotation could affect the efficiency of electron transfer process. For example, a rotation of Cyt *b*_*6*_*f* with respect to PSII could potentially speed up or slow down the diffusion and binding of the plastoquinone/plastoquinol molecular carriers in the membrane.

In conclusion, our real-time HS-AFM observation showed heterogeneous mobility of individual photosynthetic protein complexes. The photosynthetic complexes also undergo rotational diffusion, which could be an adaptive mechanism to overcome the confinement effect inherent to the thylakoid membrane. With our current HS-AFM setup, the molecular displacement of PSII and Cyt *b*_*6*_*f* was indistinguishable. Nevertheless, this characterization provides a description of the basal mobility of the photosynthetic complexes in the membrane, and the methodology can be adapted to further understand the effects of external environmental fluctuations on the global organization of the thylakoid membranes. Evidently, the successful application of HS-AFM to describe the dynamics of photosynthetic proteins in grana membranes opens a much-needed avenue to address long-standing questions regarding the dynamics of these protein complexes during photoacclimation and photoprotection mechanisms. A follow-up aim is to visualize intact thylakoids containing the complete photosynthetic electron transport chain, which will allow us to characterize molecular rearrangements induced by light fluctuations in real time.

## AUTHOR CONTRIBUTIONS

Conceived and designed the experiments: B.O., M.I., K.K.N. and C.B. Performed the experiments: B.O., S.F. and M.I. Analyzed the data: B.O. and S.F. Contributed reagents/materials/analysis tools: K.K.N. and C.B. Contributed to the writing of the manuscript: B.O., S.F., M.I., K.K.N. and C.B.

## ACKNOWLEDGMENTS

We thank Daniel Westcott and Graham Fleming for critical reading of the manuscript. This work was supported by the U.S. Department of Energy, Office of Science, through the Photosynthetic Systems program in the Office of Basic Energy Sciences. S.F. was supported in part by grant (JP15K21708) from the Ministry of Education, Culture, Sports, Science and Technology (MEXT) of Japan. K.K.N. and C.B. are investigators of the Howard Hughes Medical Institute.

## SUPPORTING MATERIAL

Nine figures and four videos are available at http://www.biophysj.org/biophysj/suppemental/

